# Harnessing Tactile Waves to Measure Skin-to-Skin Interactions

**DOI:** 10.1101/2020.05.20.105817

**Authors:** Louise P. Kirsch, Xavier E. Job, Malika Auvray, Vincent Hayward

**Author notes:** corresponding author’s.

## Abstract

Skin-to-skin touch is an essential form of tactile interaction, yet, there is no known method to quantify how we touch our own skin or someone else’s skin. Skin-to-skin touch is particularly challenging to measure objectively since interposing an instrumented sheet, no matter how thin and flexible, between the interacting skins is not an option. To fill this gap, we explored a technique that takes advantage of the propagation of vibrations from the locus of touch to pick up a signal remotely that contains information about skin-to-skin tactile interactions. These “tactile waves” were measured by an accelerometer sensor placed on the touching finger. Tactile tonicity and speed had a direct influence on measured signal power when the target of touch was the self or another person. The measurements were insensitive to changes in the location of the sensor relative to the target. Our study suggests that this method has potential for probing behaviour during skin-to-skin tactile interactions and could be a valuable technique to study social touch, self-touch, and motor-control. The method is non-invasive, easy to commission, inexpensive, and robust.

## Introduction

Skin-to-skin touch has broad implications for the sense of self (Merleau-Ponty, 1962; Crucianelli et al., 2013), body representation (Schütz-Bosbach and Haggard, 2009; van Stralen et al., 2014), affective touch (McGlone et al., 2014; Cascio et al., 2019) and motor control (Blakemore et al., 2000; Bays, 2008). It is thus connected to intriguing problems across the domains of philosophy, psychology, and neuroscience. However, to date, no empirical method is capable of measuring how we touch the skin of a living person. Even a seemingly straightforward parameter such as the tonicity of skin-to-skin touch is outside the reach of objective measurement.

When touching surfaces other than the skin, the tonicity of the motor action can be directly measured by instrumenting the touched surfaces with load sensors interposed between the surface and a mechanical reference. For example, in grasping studies, hand-held objects are typically instrumented with load cells connecting grip surfaces to the objet (e.g. (Johansson and Westling, 1984)). Such arrangements project the total interaction of the finger onto the tangential and normal directions of the touched surface. Motor behaviour can be inferred from this decomposition. Extensions of this technique using broadband sensors revealed the complexity of the fingers mechanical interactions with surfaces (Wiertlewski et al., 2011; Klöcker et al., 2013; Gueorguiev et al., 2016).

When the touched surface is the skin, it is not possible to measure the interaction by interposing an instrumented membrane between the skins since the properties of the skin contribute to the interaction (Löken and Olausson, 2010; Adams et al., 2013). As far as motor behaviour is concerned, electromyography (EMG), or acoustic myography (AMG) are invaluable techniques to investigate muscle activation (Goldenberg et al., 1991; Hodges, 2019). These techniques, however, cannot provide a precise measure of the activity of an individual at the level of the fingers, even in highly constrained conditions and with sophisticated analysis techniques (Waris et al., 2018).

Here, a novel technique is introduced which is sensitive to the effects of skin-to-skin touch and which provides a signal containing information about the behaviour of the ‘toucher’ and the nature of the interaction. It is adapted from previous work highlighting the propagation of mechanical energy in soft tissues far from a region of contact. The effect of digital tactile interactions can be measured in the whole hand (Tanaka et al., 2012; Manfredi et al., 2012; Shao et al., 2016, 2020), at least as far as in the forearm (Delhaye et al., 2012). These long-range effects are likely to result from the propagation of elastic S-waves (Vexler et al., 1999) and surface Rayleigh waves (Kirkpatrick et al., 2004) in soft tissues, with a relatively low rate of attenuation over distance.

It is known that almost all mechanical sliding contacts undergo fluctuations for any speed (Akay, 2002). The fingers are no exception. When they slide on almost any surface, including skin, contact fluctuations arise from phenomena that take place at multiple length and time scales. These phenomena vary in relative importance in accordance with the material properties of the solids in contact and the relative topographies (roughness, corrugation, conformability) at molecular, mesoscopic, and macroscopic scales (Baumberger and Caroli, 2006). The friction associated with skin-to-skin touch is the result of the skin’s complex material properties and intricate topography at all length scales. In fact, the sounds produced by the sliding of glabrous skin against glabrous skin (the ridged skin corresponding to the prehensile regions of the hand) are sufficiently strong to be heard and to modify perceptual behaviour (Jousmäki and Hari, 1998). These fluctuations are usually called frictional noise. For the present purpose they represent frictional signal.

The intensity and spectral properties of the frictional fluctuations of skin sliding against skin depend upon numerous factors, including the gross shape of the regions in contact, the type of skin, the relative states of hydration, the presence of lubricants, and of solid contaminants. Our study aimed to investigate how these fluctuations were linked to how we touch skin, including tonicity and speed. To do so, a consumer-grade accelerometer chip was attached to a single location of the touching finger to measure cutaneous vibrations remotely from the region of contact, see Fig. 1. The captured signal was compared across conditions that varied the participants’ instructed movements.

**Fig. 1.**
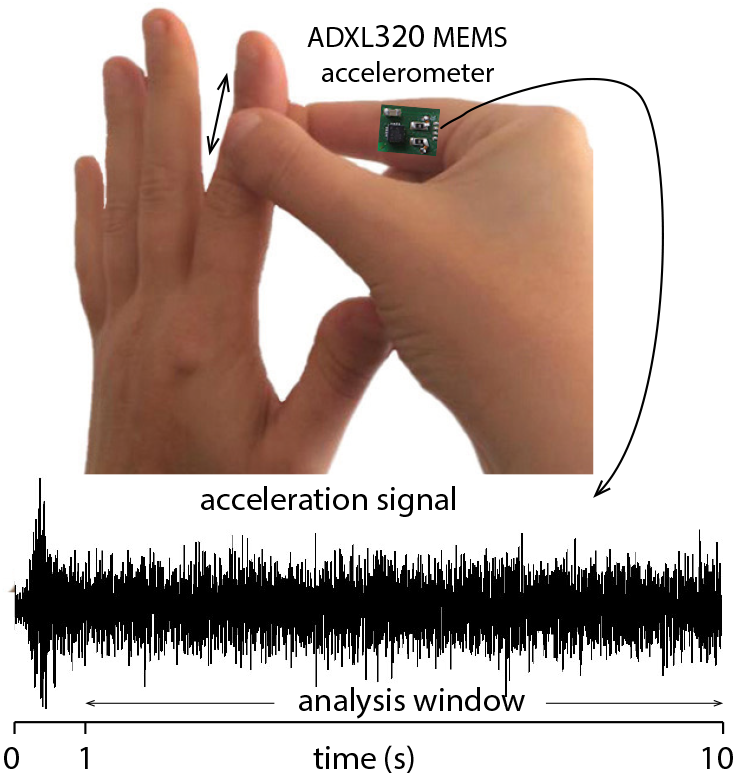
Capture of tactile waves. Signals propagated from the fingertip during tactile interaction were picked-up by consumer-grade accelerometer placed on the proximal phalange of the right index finger. The signal was acquired using a computer audio channel after 20 dB amplification.

In Experiments 1 and 2, participants were instructed to vary the tonicity of their touch (gentle or firm), or their sliding speed (fast, medium, or slow), respectively. If the signal was sensitive to these behavioural features of skin-to-skin touch, then differences in signal should be observed between these conditions (e.g. higher signal power for firm and fast compared to gentle and slow touch, respectively). In Experiment 3, the target orientation was varied such that the dorsal or ventral surface of the touched finger (i.e. the target) was facing the participant inverting the relationship of the touching fingers with the dorsal or ventral surfaces of the target. If sensor placement was critical, then the signal should depend on target orientation.

Skin-to-skin touch can be broadly divided into actions that serve to touch one’s own skin or another person’s skin, with key differences between these two types of touch (Verrillo et al., 2003; Ackerley et al., 2012). It is possible that the signal obtained during skin-to-skin touch depended on the target of the touch (e.g., (Schütz-Bosbach and Haggard, 2009). In all three experiments, the target was varied to be either the participant’s own skin, or another person’s skin in order to ascertain that the method could be applied to both types of touch.

## Experiment 1

The first experiment investigated whether the friction-induced vibration signal was sensitive to differences in the toucher’s tonicity during skin-to-skin tactile interaction. Pairs of participants touched either their own or someone else’s index finger, gently or firmly.

## Methods

### Participants

Eighteen healthy right-handed participants were recruited (ten females, mean age: 22.8 years, SD = 3.4). Participants were invited to take part in the experiment in dyads, but did not know each other. Half of the dyads were gender matched. In this and in all the experiments reported here, participants were naïve to the purpose of the experiment. Participants provided informed consent in accordance with the ethical standards outlined by the Declaration of Helsinki (1991). All experiments received approval from the university’s ethical committee. Each experiment took approximately 30 minutes to complete and the participants received payment for their participation.

### Procedure

Participants were seated opposite each other on each sides of a table approximately one meter apart. Using micropore tape, the experimenter fixed the accelerometer ventrally to the proximal phalanx of the right index finger of one of the two participants. The ‘toucher’ was then instructed to stroke her or his own left index finger (‘self’ condition) or the finger of the other participant (‘other’ condition). They used a precision grip posture such that the right index finger always touched the ventral glabrous region of the left index finger held upright, as illustrated in Fig. 2a. Participants always started the stroke from the fingertip of the target finger. One stroke consisted of one back and forth movement from the fingertip to the proximal phalanx and back.

**Fig. 2.**
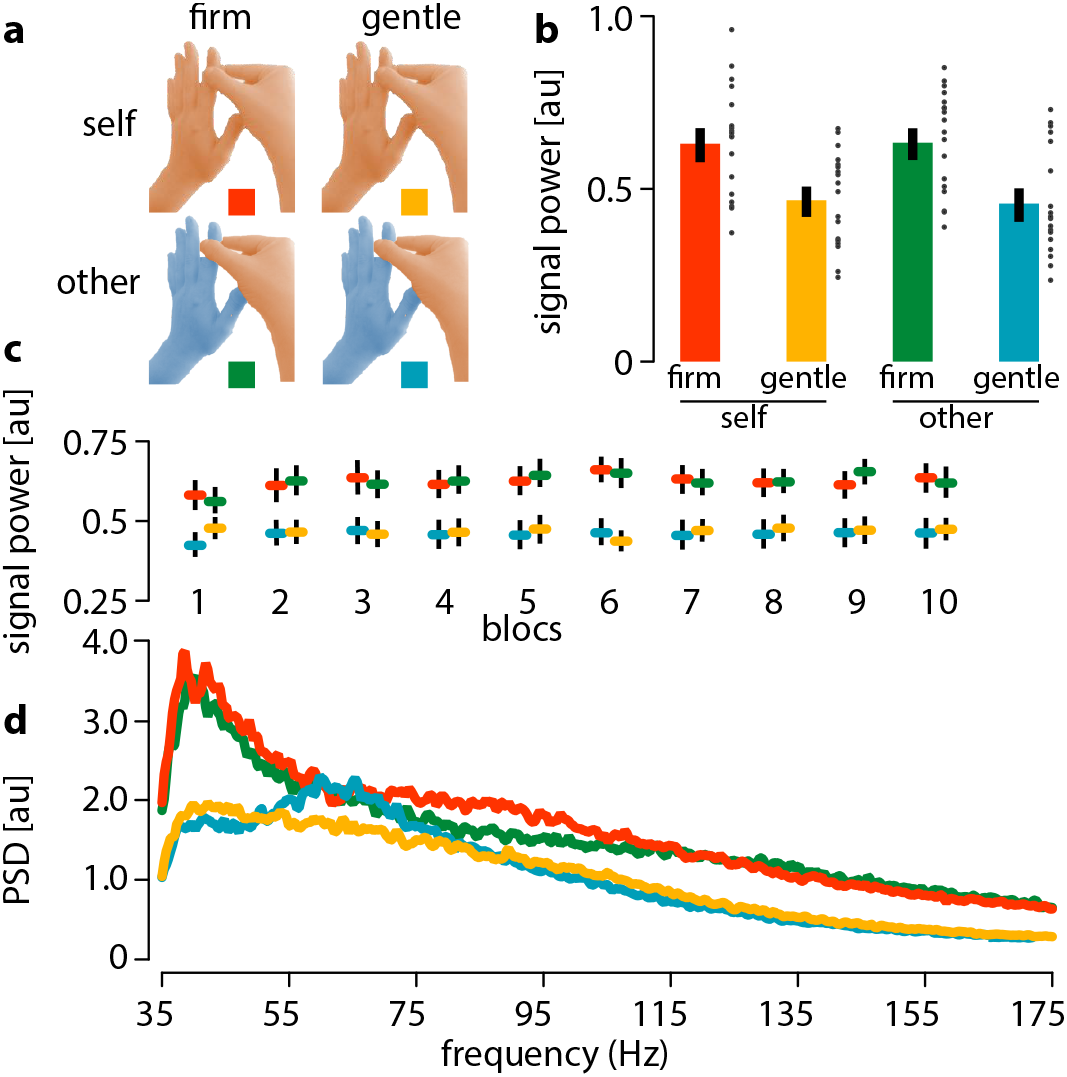
Experiment 1. **a:** Experimental design: Tonicity could be ‘firm’ or ‘gentle’, target could be ‘self’ or ‘other’; resulting in four conditions (colour coded). **b:** Total signal power of frictional fluctuations per target and tonicity conditions. Black dots show individual results. Error bars show standard error of the mean (SEM). **c:** Evolution of the average signal power by bloc number. **d:** Averaged power spectral density (PSD) over all trials and participants for each condition.

Before starting the experiment, participants completed randomised practice trials of each condition. They tried to maintain a constant pace of about one stroke per second by following a metronome (sixty beats per minutes). During the experiment, a brief sound signal (80 Hz) cued the participants to start stroking until the signal was heard again after ten seconds. Before each trial, participants were told which target to touch, their own or the other participant’s index finger, and how much to press, gently or firmly. They were free to determine what for them was gentle or firm. Each condition was randomly repeated ten times for a total of forty trials. Between each bloc, participants interchanged their places and the accelerometer was fixed to the other participant’s index finger.

### Data analysis

Only the high frequency content of the acceleration signal was considered for analysis since the low-frequency content arises from whole limb movements and changes of orientation with respect to gravity (Morris, 1973), thus mostly holding kinematic information. The first second of each trial was excluded from the analyses to eliminate the effect of the burst of signal at the transition from a static contact to a sliding contact (see Fig. 1). To minimise transducer noise, the signal was band-pass filtered in the range 35–300 Hz which is within the textural information frequency range (Wiertlewski et al., 2010). A discrete-time estimate of the average signal power was computed for each condition by assuming that the signal window was sufficiently long, a condition largely fulfilled by the audio rate sampling of 44.1 kHz. The estimates were calculated according to,

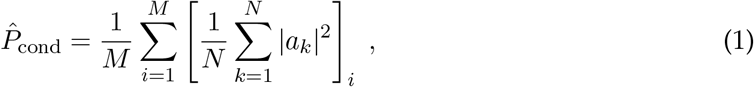

where *M* was the number of trials per condition and *N* the number of samples in the analysis window. A repeated-measure ANOVA on these averages was conducted to compare the four conditions. In addition to the analysis of signal power across the 35–300 Hz range, the power spectral density of the signal was estimated using Welch’s method to probe differences in the spectral content profiles across conditions. The analysis was applied to the averaged power spectral density of the signal in 20 Hz bands (35–55, 55–75, 75–95, 95–115, 115–135, 135–155, and 155-175 Hz). Any significant interaction was followed by post-hoc *t*-tests. All tests were Bonferroni-corrected for multiple comparisons.

## Results

A main effect of tonicity was observed (*F*(1, 17) = 32.70, *p* < 0.001, 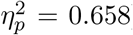); with higher signal power obtained when the touch was firm rather than gentle, see Fig. 2b. Thus, the measure was sensitive to differences in tonicity. No effect of the target nor interaction with the target were found (*F*(1, 17) = 0.009, *p* = 0.924, 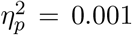, *F*(1, 17) = 0.150, *p* = 0.703, 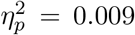, respectively). It is to note that this difference was stable over time as shown in Fig. 2c.

The difference between a gentle touch and a firm touch could also be clearly observed by inspection of the averaged power spectra over all trials and participants, see Fig. 2d, while a difference of target was not. An analysis by 20 Hz frequency bands revealed a significant effect of tonicity (*F*(1,17) = 22.616, *p* < 0.001, 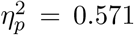), bands (*F*(1, 17) = 40.391, *p* < 0.001, 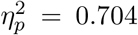), and an interaction between bands and tonicity (*F*(1, 17) = 2.942, *p* = 0.011, 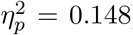). Follow-up tests showed a significant effect of tonicity for all bands above 95 Hz (all *p* < 0.001), as well as effects for the 35–55 Hz and 75–95 Hz bands (respectively: *F*(1, 17) = 9.301, *p* = 0.007, 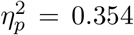 and *F*(1, 17) = 8.966, *p* = 0.008, 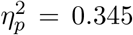), but there was no significant differences in the 55–75 Hz band (*F*(1, 17) = 1.130, *p* = 0.303, 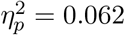).

## Experiment 2

The second experiment was designed to determine whether skin-to-skin, friction-induced vibrations were sensitive to differences in the magnitude of the sliding speed.

## Methods

### Participants

A new group of eighteen healthy right-handed individuals completed this experiment(tenfemales, meanage: 23.21 years, SD=2.55). Half of the dyads were gender matched, and gender was balanced when unmatched: half of the participants were tested by a female experimenter and the other half by a male experimenter.

### Procedure

Participants were seated to the right of the experimenter who placed the accelerometer on the participant’s right index finger. With a pen, the experimenter marked three spots on the ventral region of the participant’s left forearm, each separated by nine centimetres (creating two sites of stimulation, site 1 and site 2; see Fig.3a). These marks were identical to those made beforehand on the experimenter’s right forearm. Participants stroked with their right index finger the skin of their own forearm or that of the experimenter; alternating between site 1 and site 2, to avoid habituation. It is to note that no skin difference was expected between sites 1 and 2, so data from these two sites was averaged in the analysis. One stroke consisted of one back and forth movement between two marks. The participants synchronised their movements to a metronome set to induce three different velocities. With a 0.33 Hz beat, the average speed was low, 3.0 cm/s. At 1.0 Hz the average speed was medium, 9.0 cm/s. At 2.0 Hz, the average speed was fast, 18.0 cm/s. Each trial lasted nine seconds and each condition was repeated ten times in a randomised order, for a total of sixty trials. Participants practiced each condition for a total of six trials before data were recorded.

**Fig. 3.**
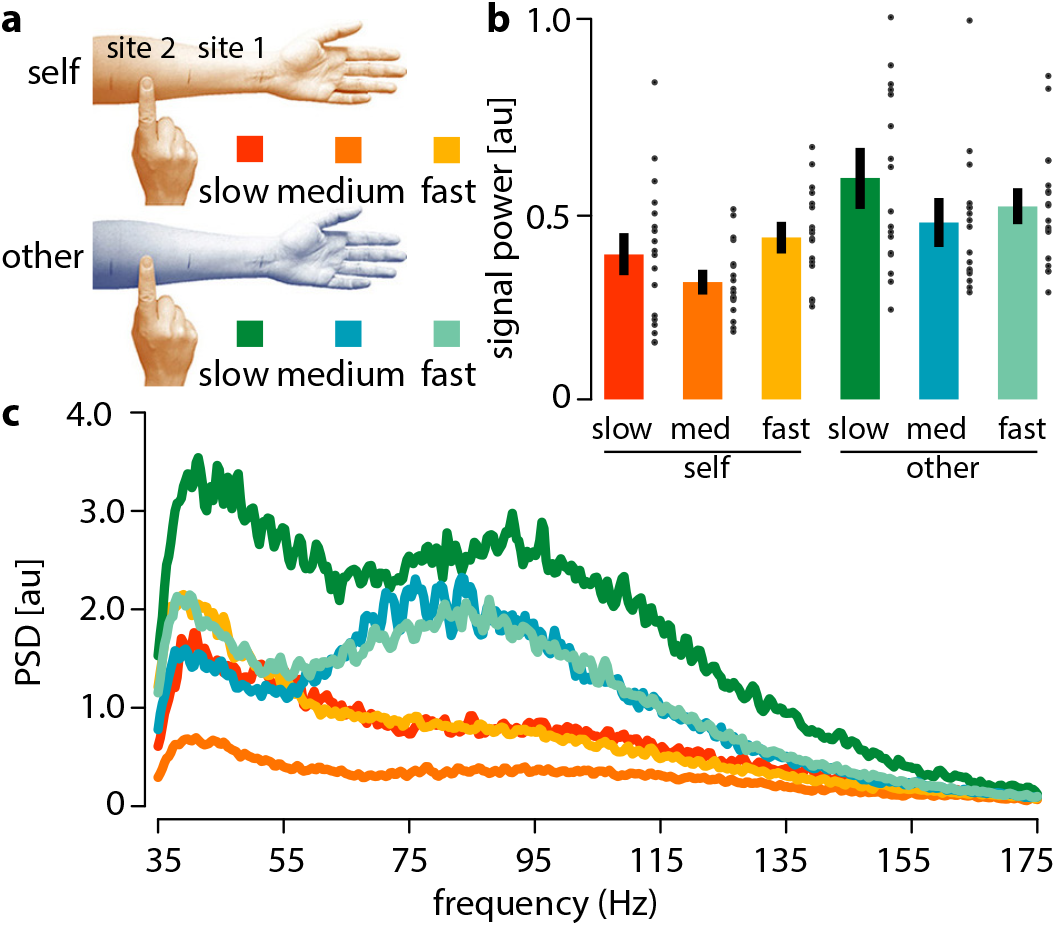
Experiment 2. **a:** Experimental design: Touching was performed at three different speeds (‘slow’, ‘medium’, ‘fast’), in random order between site 1 and 2; The target could be either ‘self’ or ‘other’; resulting in six conditions (colour coded). **b:** Total signal power of frictional fluctuations per target and speed condition. Black dots show individual results. Error bars show standard error of the mean (SEM). **c:** Averaged power spectral density (PSD) across all trials and participants over all targets and speeds.

### Results

Overall, a main effect of speed was observed (*F*(1.457, 24.768) = 6.350, *p* = 0.011, 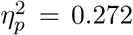) with more signal power at the highest speed, a main effect of target (*F*(1, 18) = 12.489, *p* = 0.003, 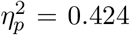), with more signal power when touching another person rather than the self, but no significant interaction (*F*(1.352, 22.980) = 2.910, *p* = 0.091, 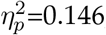; Fig. 3b). Differences between slow and medium speeds and between medium and fast speeds were found (*t*(18) = 3.042, *p* = 0.007; *t*(18) = −4.772, *p* < 0.001, respectively). The effect of target obtained here was likely due to an experimenter bias since additional analysis revealed an interaction between experimenter and target difference (*F*(1, 16) = 11.757, *p* = 0.003, 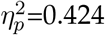) as well as a marginal main effect of target (*F*(1, 16) = 3.583, *p* = 0.077, 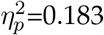); with higher power associated with one of the two experimenters (see Supplementary Fig. S1).

A frequency band analysis indicated a main effect of bands (*F*(1.846, 31.389) = 12.606, *p* < 0.001, 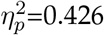), a main effect of target (*F*(1, 17) = 9.202, *p* = 0.007, 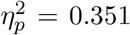), a main effect of speed (*F*(1.407, 23.913) = 5.566, *p* = 0.018, 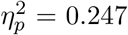), an interaction between target and bands (*F*(2.082, 35.394) = 6.176, *p* = 0.005, 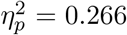), an interaction between speed and bands (*F*(2.411, 40.992) = 4.232, *p* = 0.016, 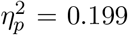) but no interaction between target and speed and no three-wayinteractionwithbands(*F*(1.972,33.521) = 1.690, *p* = 0.200, 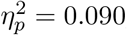, see Fig. 3c). The self-other difference was seen in the bands between 55 Hz and 155 Hz (55–75 Hz: *F*(1,17) = 11.197, *p* = 0.004, 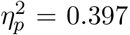; 75–95 Hz: *F*(1, 17) = 11.993, *p* = 0.003, 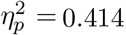; 95-115 Hz: *F*(1, 17) = 10.635, *p* = 0.005, 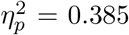). However, the effect of speed and the interaction between speed and target did not survive Bonferonni correction (*α* = 0.0074) in any of the bands.

## Experiment 3

Experiment 3 investigated whether the signal varied with the orientation of the target hand since the location of the sensor relative to target may have influenced the signal.

## Methods

### Participants

A new group of eighteen right-handed participants completed the experiment (nine females, mean age: 23.6 years, SD = 3.6). Participants were invited to take part in the experiment in dyads, but they did not know each other. As in Experiments 1 and 2 gender was balanced across dyads.

### Procedure

As in Experiment 1, participants were seated opposite each other on either side of a table approximately one meter apart. The accelerometer was placed on the right index finger of one participant of the dyad, who would be the participant performing the touch. The accelerometer was fixed in the same position as in Experiments 1 and 2, thus distance between the sensor and the regions of contact varied with target orientation. Participants performed the same action as in Experiment 1 (precision grip), with the sole difference being the orientation of the touched index finger (i.e. target orientation; see Fig 4a). In the ‘outwards’ condition, the palm of the target hand faced away the toucher (i.e. the active index of the toucher was in contact with the glabrous skin on the ventral side of the target finger, and the thumb with the hairy skin on the dorsal side). In the ‘inwards’ condition, the palm of the target hand faced towards the toucher (i.e. the reversed configuration). As in Experiment 1 and 2, the ‘toucher’ was instructed to stroke their own left index finger (‘self’ condition) or the finger of the other participant (‘other’ condition). No tonicity instruction was given. Participants were encouraged to keep a constant speed by the same method as in Experiment 1. Each condition was repeated ten times for a total of forty randomised trials. After those trials, the two participants interchanged places and the accelerometer was attached to the new toucher.

**Fig. 4.**
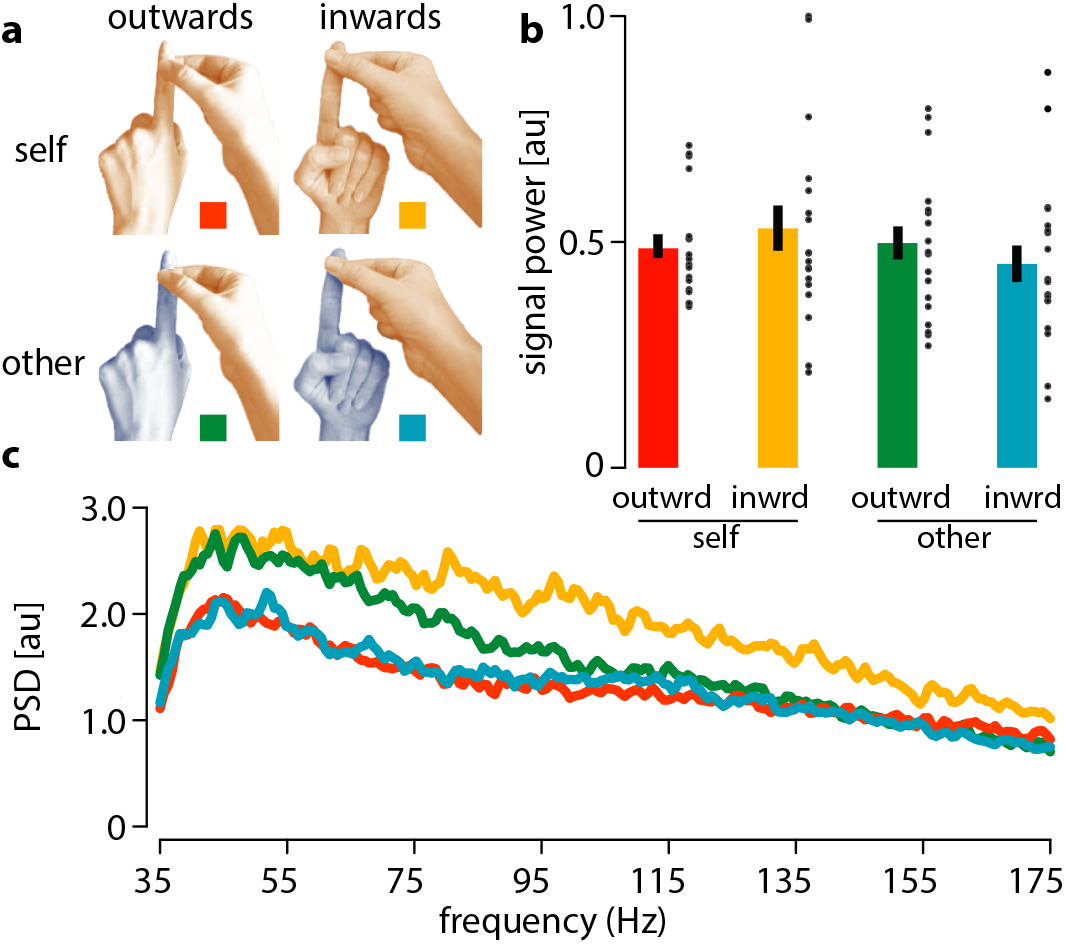
Experiment 3. **a:** Experimental design: Target orientation could be ‘outwards’ or ‘inwards’, target could be either ‘self’ or ‘other’; resulting in four conditions (colour coded). **b:** Total signal power of frictional fluctuations per target and target orientation; Black dots show individual results. Error bars show standard error of the mean (SEM). **c:** Averaged power spectral density across all trials for each condition.

## Results

The results showed no effect of target and no effect of target orientation (target: *F*(1, 17) = 1.724, *p* = 0.207, 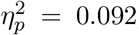; target orientation: *F*(1, 17) = 0.002, *p* = 0.969, 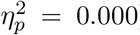), but they showed an interaction between target and skin type (*F*(1, 17) = 6.393, *p* = 0.022, 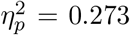), see Fig. 4b. However, none of the post-hoc *t*-test survived Bonferroni correction, suggesting no significant impact of the orientation of the target.

An analysis by frequency bands, Fig. 4c, revealed a main effect of bands (*F*(1.513, 25.725) = 27.472, *p* < 0.001, 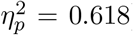), an interaction between target and skin type (*F*(1, 17) = 5.239, *p* = 0.035, 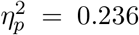), and a three-way interaction with bands (*F*(1.815, 30.860) = 5.221, *p* = 0.013, 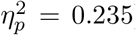). Follow-up analyses did not yield to any significant results (no main effect of target, skin nor interaction survived the Bonferroni correction).

## Discussion

Skin-to-skin touch is challenging to measure objectively, yet it presents a number of intriguing problems that span neuroscience, psychology and philosophy. Here, we tested the efficacy of a new measure of skin-to-skin tactile behaviour that took advantage of the frictional fluctuations propagating in soft tissues (Shao et al., 2016, 2020). Participants were instructed to stroke skin surfaces while an accelerometer was fixed to their touching finger. The recorded signal contained information about the vibrations elicited during touch. Participants varied the tonicity of their touch, their movement speed, the orientation of the target, as well as the target identity (self-touch vs. touching another’s skin).

The analysis relied on the total signal power and the distribution of this power in specific frequency bands. The signal exhibited considerable variability between individuals, however this limitation is shared by most other physiological signal measurements including pupil dilation, e.g. (Einhäuser et al., 2008; Wierda et al., 2012), skin conductance, e.g. (Tronstad et al., 2010; van Dooren et al., 2012), electromyography, e.g. (Goldenberg et al., 1991), respiration, e.g. (Boiten et al., 1994; Valderas et al., 2015) and heart-rate, e.g. (Appelhans and Luecken, 2006; Garfinkel et al., 2015). Despite high inter-individual variability, useful information could be extracted from the signal, allowing comparisons across experimental conditions.

Experiment 1 showed a clear effect of touch tonicity when participants were instructed to apply either gentle or firm pressure. The signal power was significantly higher during firm compared to gentle touch. This demonstrates that a consumer-grade accelerometer is able to capture tactile signals and can be used as a proxy of the force applied during skin-to-skin touch. Therefore, the method is able to detect differences in the tonicity of skin-to-skin touch.

Experiment 2 showed that the signal was sensitive to the speed with which participants touched the skin. The relationship between sliding speed and signal power was however complex. The medium speed (9 cm/s) elicited significantly lower signal power than the faster speed (18 cm/s) and the slower speed (3 cm/s). There may be several reasons why the relationship between movement speed and signal power was not monotonic. Participants probably moved less smoothly at slower speeds (Guigon et al., 2019). Jerky movements may have caused bursts of signal at the slowest speed. The observation of greater signal power at the highest speed (18 cm/s) is in line with our initial hypothesis since greater frictional energy was dissipated during the same time window.

The positioning of a single sensor relative to the source of contact may have had an effect on the signal obtained, particularly with differences across experimental conditions. Experiment 3 assessed the influence of target orientation on the signal obtained during skin-to-skin touch. In Experiment 1, participants gripped the finger when it was oriented with the dorsal surface facing towards them. In Experiment 3, the target orientation was manipulated to either be the same, as in Experiments 1, or oriented with the ventral surface facing toward the toucher. The signal power did not vary systematically with target orientation, suggesting that a similar signal would have been obtained from a sensor placed on the active thumb rather than active index finger. In practice, this means that experimenters are not constrained by specific placements of the sensor on the hand.

Several lines of evidence suggest that we may touch ourselves differently from others, this is the case, for example in the “touchant-touché” phenomenon (Husserl, 1989; Merleau-Ponty, 1962; Schütz-Bosbach and Haggard, 2009). The literature also suggests that self-generated touch is perceived to be less intense than externally generated touch (Blakemore et al., 2000; Shergill et al., 2003; Bays, 2008). In Experiments 1 and 3, participants touched themselves or another person in dyads. The target had no influence on signal power. In Experiment 2, one of two experimenters was the ‘other’ target. Stronger signal power was found when participants touched another person. Further analyses revealed that the signal was higher with one of the two experimenters. Overall, our results did not show clear differences between touching one’s own skin compared to another person’s skin. This finding may seem surprising given the known differences between touch applied to one’s own compared to another person’s skin (Verrillo et al., 2003; Ackerley et al., 2012). However, the lack of difference may reveal the existence of a robust motor invariant that is insensitive to the target of touch, particularly under the conditions of Experiment 1. Several motor invariants related to motor tonicity have been documented Feldman (1980); Latash et al. (2007). In Experiment 2, having only two ‘other’ targets may have reduced variability and introduced additional factors such as skin hydration and also possible gender effects (that were balanced in Experiment 1 and 3, as shown in Supplementary Fig. S1). This result suggests that our method could be applied to differentiate between targets. Future studies could investigate the relative advantages of various stroking actions to extract specific types of information from the vibration signal.

Our results were obtained using spectral density analyses, including total signal power and power spectral density in broad frequency bands. However, in natural touch, cutaneous vibrations are almost always non-stationary signals, which means that the generating processes varies over time. In our study, power spectral density analyses were adequate for the investigated factors because the participants were instructed to repeat the same action over relatively long periods of time. Future research based on the analysis of time-varying phenomena could certainly be possible, for example, with short-time Fourier analysis.

Future research may be also be aimed at estimating the source of touch, or even the type of action executed, from vibrations signals measured in the hands. Blind source separation analysis techniques (Comon and Jutten, 2010) could be used since the frictional fluctuations come from sources arising from phenomena associated to different length scales. Another direction would be to increase the number of accelerometer sensors across the hand as in Shao et al. (2016, 2020) who used up to thirty sensors. Finally, an abundance of tools based on machine learning techniques are now available that are able to extract information from complex signals. Such methods could be used to decode behavioural interactions from the resulting tactile vibrations.

To conclude, the results demonstrated the direct measurement of cutaneous vibrations resulting from friction elicited by skin-to-skin contact. We showed that the signal is primarily sensitive to the tonicity and the speed of tactile interactions. The measure has significant potential for probing behaviour during skin-to-skin tactile interactions, opening avenues for future research investigating a variety of factors underlying self-touch as well as social touch and motor control.

## Acknowledgements

We thank Agnès Roby-Bramy for insightful discussions. This work was supported by Agence nationale de la recherche grant ANR-16-CE28-0015 “Developmental Tool Mastery” led by Alessandro Farné. The authors declare that no competing interests exist.

## Open practice statement

The data for all experiments are available on the OSF repository and can be accessed via this link: https://osf.io/7gw5z/?view_only=7d351d4a7b6a443392157da6bb643a90

## Supplementary Information

Half of the participants in Experiment 2 were tested with a male experimenter as target and the other half with a female experimenter. Higher signal power was observed when the participants touched the male experimenter compared to the female experimenter. Gender did not influence how they touched their own forearm.

**Fig. S1.**
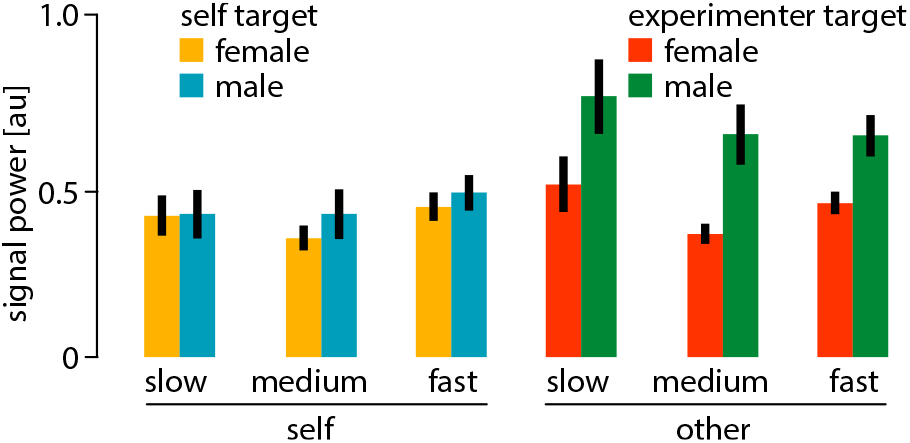
Experimenter effect in Experiment 2.

